# Quantum algorithm for identifying RNA 3D motifs by processing RNA secondary structure graphs

**DOI:** 10.1101/2025.04.25.650554

**Authors:** Jing-Kai Fang, Fengmin Guo, Teng Ma, Yong Zhang, Ou Wang, Jun-Han Huang, Yuxiang Li

**Affiliations:** BGI Research, Shenzhen 518083, China; BGI Research, Beijing 102601, China; State Key Laboratory of Genome and Multi-omics Technologies, BGI Research, Shenzhen 518083, China; BGI Research, Wuhan 430047, China; Guangdong Bigdata Engineering Technology Research Center for Life Sciences, BGI Research, Shenzhen 518083, China

## Abstract

The identification of RNA motifs plays a crucial role in understanding biological functions, predicting RNA 3D structures, and designing RNA-based therapeutics. One of the efficient strategies approaches this task by analyzing secondary structure graphs annotated with non-canonical base interactions, effectively reframing the problem as a two-dimensional graph-theoretic challenge. However, both the graph isomorphism problem and the maximum common subgraph problem in this domain are classified as NP-complete. Classical computing architectures struggle to address these problems efficiently due to their substantial computational resource demands. In this work, we propose a quantum algorithm specifically designed for processing RNA secondary structure graphs. By encoding the adjacency matrix into a quantum circuit and incorporating base interaction families as edge weights, our approach achieves a resource efficiency of 𝒪**(log**_**2**_ ***N***) in terms of qubits. Numerical simulations demonstrate the algorithm’s capability to identify 15 distinct motifs across three structural categories: hairpin loops, internal loops, and three-way junction loops. Furthermore, the method successfully detects the common motif between two complete RNA chains. Finally, 29 high conserved RNA motifs were identified in four cell lines of a key lung cancer biomarker gene (MIF), and their potential roles as drug targets were explored. This work presents a highly promising quantum computing framework for addressing the RNA motif identification problem.

## 1 Introduction

RNA motifs, defined as recurrent structural elements in RNA molecules, play a crucial role in both the molecular structure stability and biological function. Characterized by specific arrangements of non-canonical base pairs, these motifs are exhibit complex three-dimensional conformations that extend beyond the planar configurations defined by canonical Watson-Crick and Wobble base pairs [1]. Representative motifs such as GNRA loops [2], Kink-turns [3], G-bulges [4], and A-minors [5], demonstrate high evolutionary conservation across diverse RNA molecules, reflecting selective pressures to maintain specific interaction patterns [6]. These patterns impose constraints on sequences for proper folding, thereby establishing a correspondence between nucleotide sequences and their three-dimensional configurations.

Identifying and characterizing RNA motifs are essential for understanding RNA evolution and developing predictive models of RNA three-dimensional structures, which provide insights into how RNA molecules fold [7–10] and interact with other biomolecules. This understanding is crucial for studying the evolutionary conservation of RNA structures, designing synthetic RNAs with specific functions [11, 12] and developing small-molecule drugs that target RNA [13–16] in the fields like synthetic biology and therapeutic development. Furthermore, recognizing RNA motifs also aids in elucidating the mechanisms of RNA-mediated gene regulation [17–19] and the overall architecture of complex RNA molecules.

Specialized databases such as the RNA 3D Motif Atlas [20] and RNA Bricks [21] systematically catalog these structural motifs, enabling applications in synthetic biology and RNA engineering through comprehensive annotations of three-dimensional substructures. These resources further accelerate the discovery of novel RNA-protein interactions and RNA-ligand binding patterns, providing critical foundations for therapeutic development and mechanistic studies in molecular biology. Computational approaches employ two distinct paradigms for motif detection: (1) sequence-driven methodologies exemplified by MEME [22], QRNA [23], RNAz [24], and CMfinder [25], which leverage evolutionary conservation patterns; and (2) structure-centric frameworks such as RNAMotifScan [26], FR3D [27] and so on [28–38] that utilize graph-theoretic representations of RNA architecture. These methodology ranges from recognition of known motifs to de novo identification of structural modules through geometric pattern detection. The synergistic integration of these computational tools with structural databases facilitates the advancements in multidimensional investigations of RNA motifs mentioned above.

The application of quantum algorithms to RNA-related research [39–41] is gaining significant traction, driven by inherent limitations in classical computational architectures when processing rapidly growing sequencing datas. With classical supercomputers approaching fundamental physical constraints dictated by Moore’s law, the scientific community urgently requires post-Moore computational frameworks. Quantum computing offers a promising solution with its enhanced computational efficiency and ability to tackle complex optimization problems [42, 43]. Studies have demonstrated that quantum algorithms can provide potential advantages in various bioinformatics tasks, including genome assembly [44–48], sequence alignment [49–51], protein folding [52–54], and phylogenetic tree inference [55]. These advancements have sparked interest in exploring quantum computing for RNA-related challenges, such as predicting RNA tertiary structures and identifying RNA motifs. By leveraging quantum algorithms, computational bottlenecks faced by classical methods may be overcome, enabling more accurate and efficient RNA data analysis. This progress could deepen our understanding of RNA biology and contribute to the development of novel RNA-based therapeutics and synthetic biology applications.

In this work, we present a quantum algorithm for RNA motif identification and demonstrate how RNA secondary structures can be effectively encoded into quantum circuits. The algorithm successfully identifies known RNA motifs and common motifs between two RNA sequences using real data. Furthermore, we apply our method to analyze the secondary structures of four lung cancer cell lines, identifying conserved motifs across all samples. These findings offer valuable insights for the design of RNA-targeted small-molecule therapeutics.

## 2 Results

### Dataset

A non-redundant RNA database of RNA3DHud [56] with version 3.87 (August 14, 2024) is used in this work. It comprises a non-redundant collection of recurrent RNA 3D hairpin, internal and three-way junction loop motifs, along with the structural annotations of the corresponding RNAs. As shown in Fig. 2, we extracted five RNA motifs of varying sizes from each motif class and provided their RNA secondary structure graphs with interaction annotations. Note that for RNA motifs, a node in the secondary structure graph corresponds to one of the four nucleotide types. Therefore a set of abbreviations as shown in Table 1 is used to indicate all possible nucleotides allowed for each node. In this work, we annotate only base pair interactions, including both canonical and non-canonical base pairs, while excluding base stacking and base-backbone interactions. This exclusion is because we only focus on identifying similar substructures in RNA 3D structures, as complex interactions may reduce the generalizability of the substructures. Full interaction annotations can be utilized for more nuanced alignment and analysis tasks, which are beyond the scope of this work. And the 3D structure of RNAs are obtained from PDB [57] database for motif identification. Furthermore, identification of conserved motifs is perfromed to four lung cancer cell lines: NCI-H23, NCI-H2030, NCI-H1793 and NCI-H358, which come from smartSHAPE technology.

**Table 1.** Abbreviations for nucleotide types. RNA motifs can adopt similar three-dimensional structures from diverse base sequences; consequently, secondarystructure graphs employ nucleotidetype abbreviations to concisely represent all allowable bases at each position.

| Abbreviation | Nucleotide type | Abbreviation | Nucleotide type |
| --- | --- | --- | --- |
| Y | pyrimidines(C or U) | D | not C |
| R | purines(G or A) | H | not G |
| W | weak(A or U) | A | only A |
| S | strong(G or C) | U | only U |
| K | keto(G or U) | C | only C |
| M | amino(A or C) | G | only G |
| B | not A | N | any nucleotide |
| V | not U |  |  |

### RNA 2D structure graphs

Non-canonical base pairing plays a key role in RNA folding to form unique three-dimensional structures. An RNA motif can be defined as a specific sequence or structural pattern that folds into essentially identical three-dimensional structures, characterized by specific arrangements of isosteric non-canonical base pairs within RNA. To identify and compare RNA motifs, RNA 2D structure graphs with base interaction annotations can be used without losing essential three-dimensional structural information. In these graphs, each node represents a nucleotide, and each edge represents an interaction between nucleotides, including base pairing and the phosphodiester backbone, distinguished by edge weights. The annotations of base pair interactions adhere to the Leontis-Westhof geometric classification [58]. This classification system categorizes interactions into 12 distinct geometric families, based on the intricate hydrogen bond interactions that occur between RNA bases. These interactions take place on three specific edges of an RNA base: the Watson-Crick edge, the Hoogsteen edge, and the Sugar edge. Additionally, each edge of one base may interact with an edge of another base in either the cis or trans orientation of the glycosidic bond. Thus, as shown in Table 2, non-canonical base-pair interactions are annotated with symbols representing the type and relative orientation of the interacting edges.

**Table 2.**
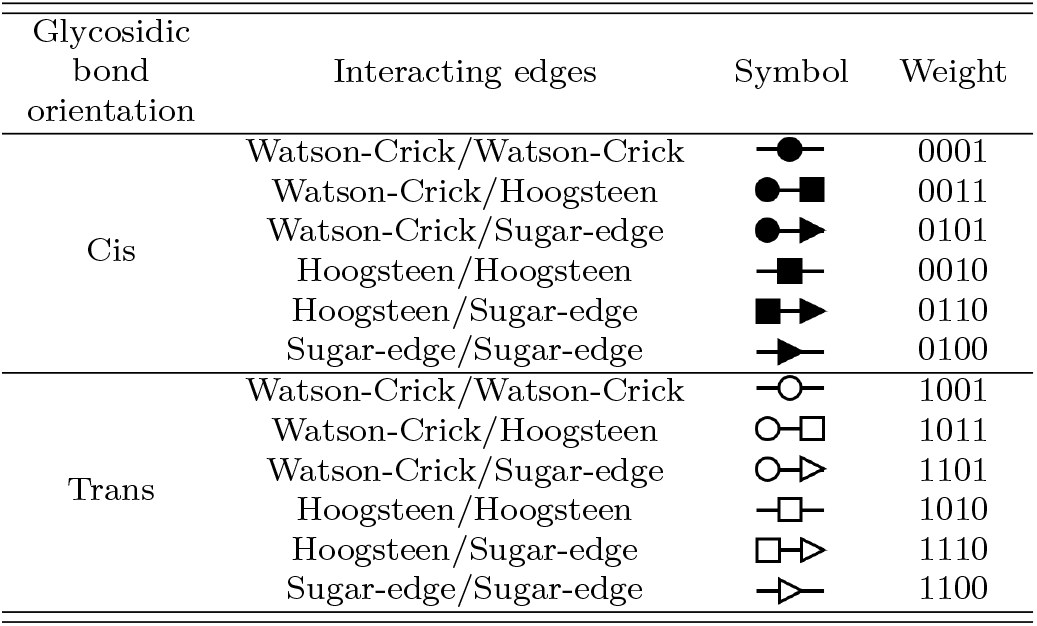
12 geometric families of base interactions and their corresponding symbols. Their corresponding binary weights are also indicated.

### Binarization of the adjacency matrix

For an RNA 2D structure graph *G* = {*V, E*}, each vertex *V* corresponds to the base type of a nucleotide, and each edge *E* represents either a phosphate backbone connection or a basepair interaction. The annotations can be easily converted into weights corresponding to each edges, requiring only 13 discrete values. A 4-bits binary number is sufficient to represent the weights of all edges, expressed as 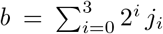, where *j*_*i*_ ∈ {0, 1} denotes the value of the *i*^*th*^ binary bit. Here we focus on implementing the binary representation of the weights associated with edges annotations in the RNA secondary structure graph. The reason for using binary representation is discussed in Section 4. While it is possible to randomly assign a 4-bit binary weight to different types of interactions, we believe that considering the similarity relationships between different edges can lead to better results, particularly in cases of imperfect matching. Therefore, we take into account the differences in each bit between two distinct binary weight values. Specifically, we select 13 binary weights out of 16 to maximize the summation of XORs of pairs of binary weights corresponding to different interacting edges, as follows:

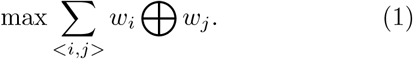

The corresponding weights of basepair interactions are presented in the last column in Table 2. And we assign a weight of 1000 to the phosphatebackbone edges to maximize their distinction from basepair interactions. By converting each matrix element into a 2×2 block and filling the blocks with the corresponding binary weights, the adjacency matrix can be transformed to a binary matrix. As shown in Fig. 1(a), the original weighted *N* × *N* adjacency matrix of RNA secondary structure graph is expanded into a 2*N* × 2*N* binary adjacency matrix.

**Fig. 1.**
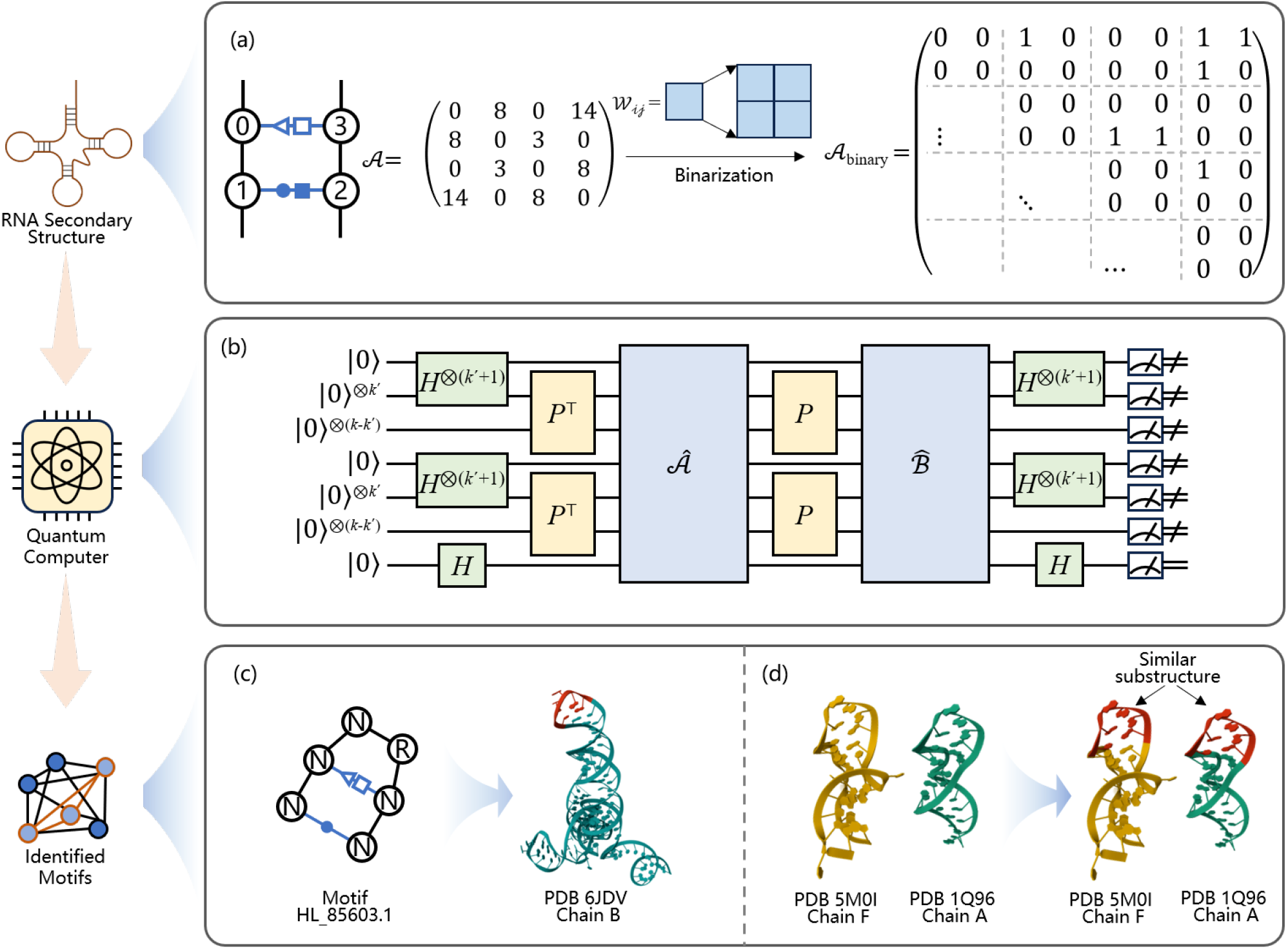
Schematic representation of the quantum algorithm for RNA motif identification. (a) Binarization process for an adjacency matrix of RNA secondary structure graph with base pair interations annotation, (b) The circuit of subgraph isomorphism algorithm with labeled edges. Here *k*(*k*^*′*^) = log_2_[*N* (*N*^*′*^)] with *N* (*N*^*′*^) being the number of vertices of the graph (subgraph), (c) Identification of motif HL_85603.1 in PDB 6JDV chain B and (d) Identification for two RNA chains: PDB 5M0I chain F(Secquence:GAUAACUGAAUCGAAAGACAUUAUCACG) and PDB 1Q96 chain A(Secquence:GGUGCUCAGUAUGAGAAGAACCGCACC). They share an identical motif HL_85603.1, which is a 6-nucleotide hairpin loop possessing a Hoogsteen/Sugar edge base pair and a Watson Crick base pair.

**Fig. 2.**
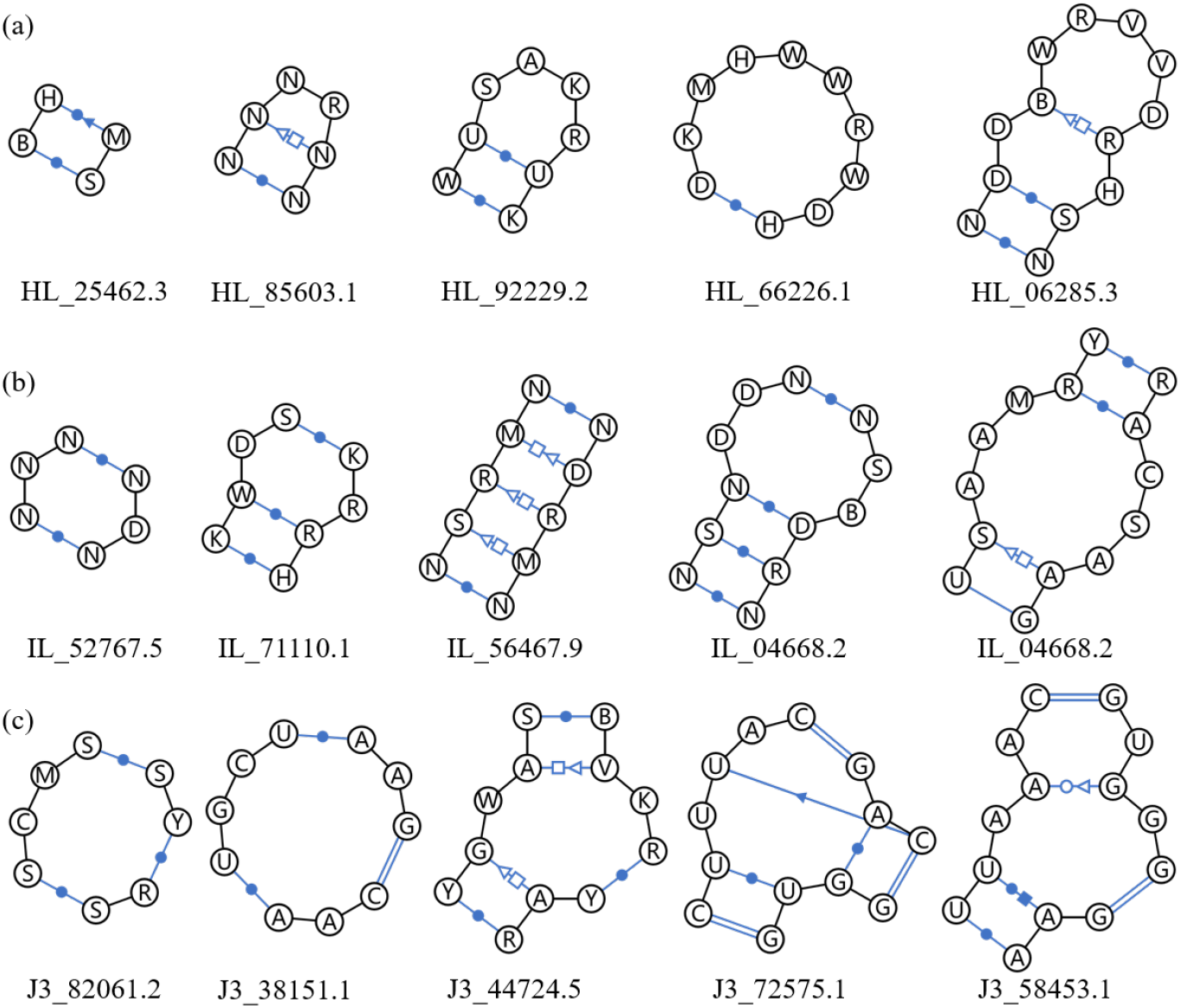
Diagram of the secondary structure of RNA motifs with five different sizes within (a) Hairpin loop, (b) Internal loop, and (c) 3-way Junction loop, utilized RNA motif identification. Each motif is annotated with base interactions containing non-canonical base pairs. And the posssible nucleotide types for them are also annotated.

**Fig. 3.**
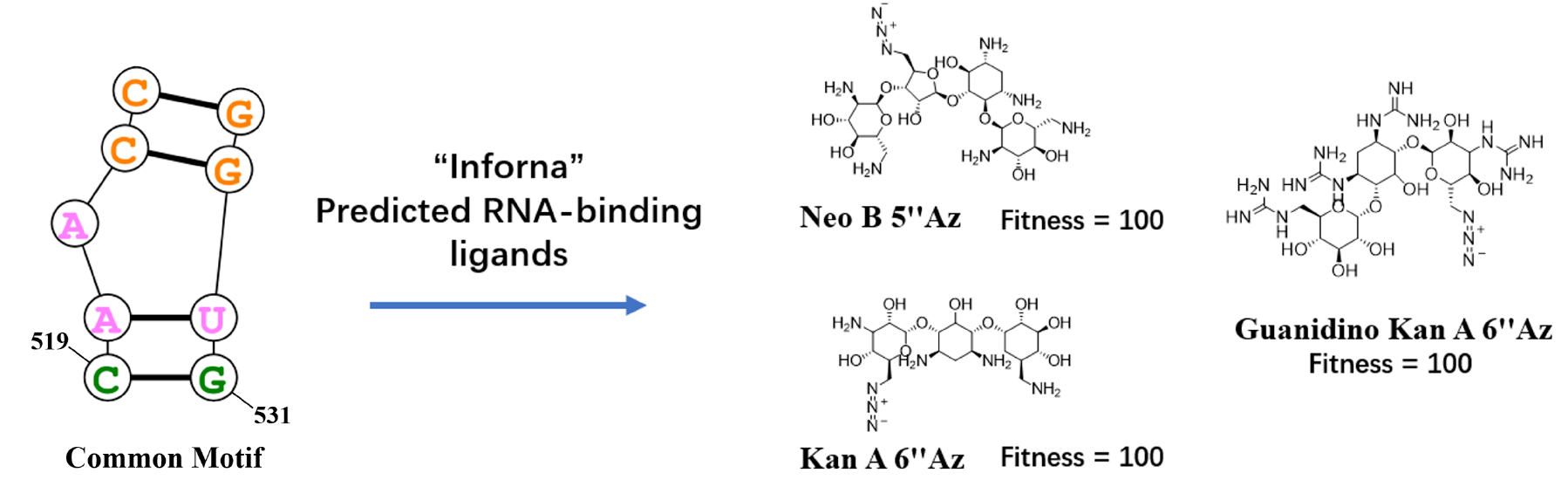
This schematic illustrates the prediction of RNA-targeted small-molecule drugs: conserved motifs identified in lung cancer cell lines by our quantum algorithm are submitted to the Inforna platform to generate candidate therapeutics.

Moreover, the diagonal entries of the adjacency matrix can be used to represent intrinsic properties of the nodes. In this context, each 2×2 diagonal block precisely encodes the four RNA base types as follows: {*A* →1000, *U* → 0100, *C* →0010, *G* →0001}. This representation can also accommodate cases where multiple base types are allowed by using the sum of the corresponding binary values. Such flexibility is particularly useful for RNA motifs that permit different sequences to adopt similar three-dimensional structures.

### Identification of RNA motif

We analyze five motifs of varying sizes—each less than 16 nucleotides (nt) in length—across three motif families: hairpin loops, internal loops, and 3-way junction loops. Motif identification is then performed to determine the precise location of each motif on the RNA chain. Due to the computational limitations of quantum simulator when processing long RNA sequences, we truncate real RNA chains. Specifically, we select a 16-nt segment to ensure that it contains the previously identified motifs within its boundaries.

Since we do not impose restrictions on the direction of interactions, the resulting RNA secondary structure is represented as an undirected graph. However, we limit the types of nucleotides and partially relax the identification constraints. While ensuring that the optimal solution can be identified, we relax the constraint on interaction direction to some extent. This adjustment improves optimization performance and simultaneously allows for the inclusion of suboptimal solutions with incomplete matching that may still hold biological significance.

As shown in Table 3, we perform 100 experiments for each motif identification task. Except for the smallest motif with a length of 4, all motifs demonstrated a fidelity exceeding 90%. The lower fidelity for smaller motifs is attributed to the substantial relative differences between suboptimal and exact solutions. For instance, in four-nucleotide motifs, a single nucleotide discrepancy results in a 25% deviation. However, nearest neighbor search optimization effectively resolves these suboptimal cases, enabling the identification of the exact solution. Notably, there is no significant difference in the number of exact solutions ultimately identified between smaller and larger motifs. As the motif size increases, the number of exact solutions fluctuates within a specific range, without showing a clear correlation with motif size.

**Table 3.** Statistical table for the identification tasks of three types of motifs: hairpin loops, internal loops, and 3-way juntion loops. 100 identifications were performed for each motif using our algorithm.

| Motif Type | Size(nt) | Search RNA(PDB) | Avg Loss | Avg Iteration | Exact Solution |
| --- | --- | --- | --- | --- | --- |
| Hairpin Loop | 4 | 4WT8 | -0.851175 | 66.1 | 70 |
|  | 6 | 6JDV | -0.926060 | 90.4 | 80 |
|  | 8 | 5TBW | -0.923589 | 68.1 | 70 |
|  | 10 | 2A64 | -0.930637 | 82.7 | 70 |
|  | 13 | 4WFL | -0.952204 | 69.3 | 85 |
| Internal Loop | 6 | 2ZI0 | -0.936874 | 86.5 | 79 |
|  | 8 | 1KUQ | -0.946855 | 62.3 | 52 |
|  | 10 | 3IGI | -0.937561 | 76.4 | 65 |
|  | 12 | 6CZR | -0.9620315 | 67.0 | 89 |
|  | 14 | 4V9F | -0.952192 | 62.4 | 92 |
| 3-Way Junction Loop | 8 | 5T5A | -0.933445 | 67.9 | 67 |
|  | 10 | 6ZDP | -0.954188 | 69.6 | 93 |
|  | 12 | 8VTW | -0.948444 | 61.2 | 93 |
|  | 13 | 3NDB | -0.949851 | 64.2 | 78 |
|  | 14 | 6JQ5 | -0.950186 | 68.8 | 85 |

### Identification of common substructures in two RNA chains

Beyond identifying specific RNA motifs within a single RNA chain, our algorithm also identifies similar substructures across different RNA sequences. This task facilitates the discovery of novel RNA motifs and enables the analysis of structural similarities among diverse RNAs. Such insights are invaluable for exploring the relationship between conserved RNA three-dimensional structures and their associated biological functions.

We performed motif identification on two complete RNA chains: PDB 5M0I chain F and PDB 1Q96 Chain A, which share a common substructure, Motif HL_85603.1. When aligning these RNA chains, we omit base type encoding and focus solely on structural similarity, as similar RNA 3D substructures can often arise from different nucleotide sequences. Upon completing the alignment using our quantum algorithm, we obtain a permutation matrix *P*. As described in Section 4, it represents the correspondence of each vertex in two RNA secondary structure graphs. Then, an additional post-processing step is applied to remove points with unmatched edges, allowing for the extraction of perfectly matched substructures. As shown in Fig. 1(d), our method successfully identifies a common hairpin loop substructure, which corresponds exactly to Motif HL_85603.1.

### Identification of RNA motifs and discovery of potential small-molecule binders

The identification of RNA motifs and their potential small-molecule binders not only provides a critical theoretical foundation for the development of RNA-targeted drugs but also establishes an essential basis for optimizing novel therapeutic strategies such as RlBOTACs (Ribonuclease Targeting Chimeras). In recent years, advances in RNA biology yield growing evidence that specific functional motifs within RNA secondary structures serve as highly selective binding sites for small molecules, enabling precise modulation of gene expression or targeted degradation [16].

In this study, we innovatively employ an efficient quantum computing-based algorithm that significantly improves the identification efficiency of RNA structural motifs. Utilizing smartSHAPE technology, we obtain transcriptome-wide RNA secondary structures from four lung cancer cell lines (NCl-H23, NCI-H2030, NCl-H1793, and NCl-H358; transcriptomic features are provided in Fig. 4. Taking Macrophage Migration Inhibitory Factor (MIF), a key lung cancer biomarker gene, as an example, our quantum algorithm systematically analyzes the similarities and differences in RNA structural motifs across the four cell lines. This analysis leads to the identification of 29 highly conserved RNA motifs (with structural distribution and sequence conservation analysis shown in Fig. 5. The conservation of these motifs across all cell lines suggests their structural stability and a potential critical role in MIF gene regulation.

**Fig. 4.**
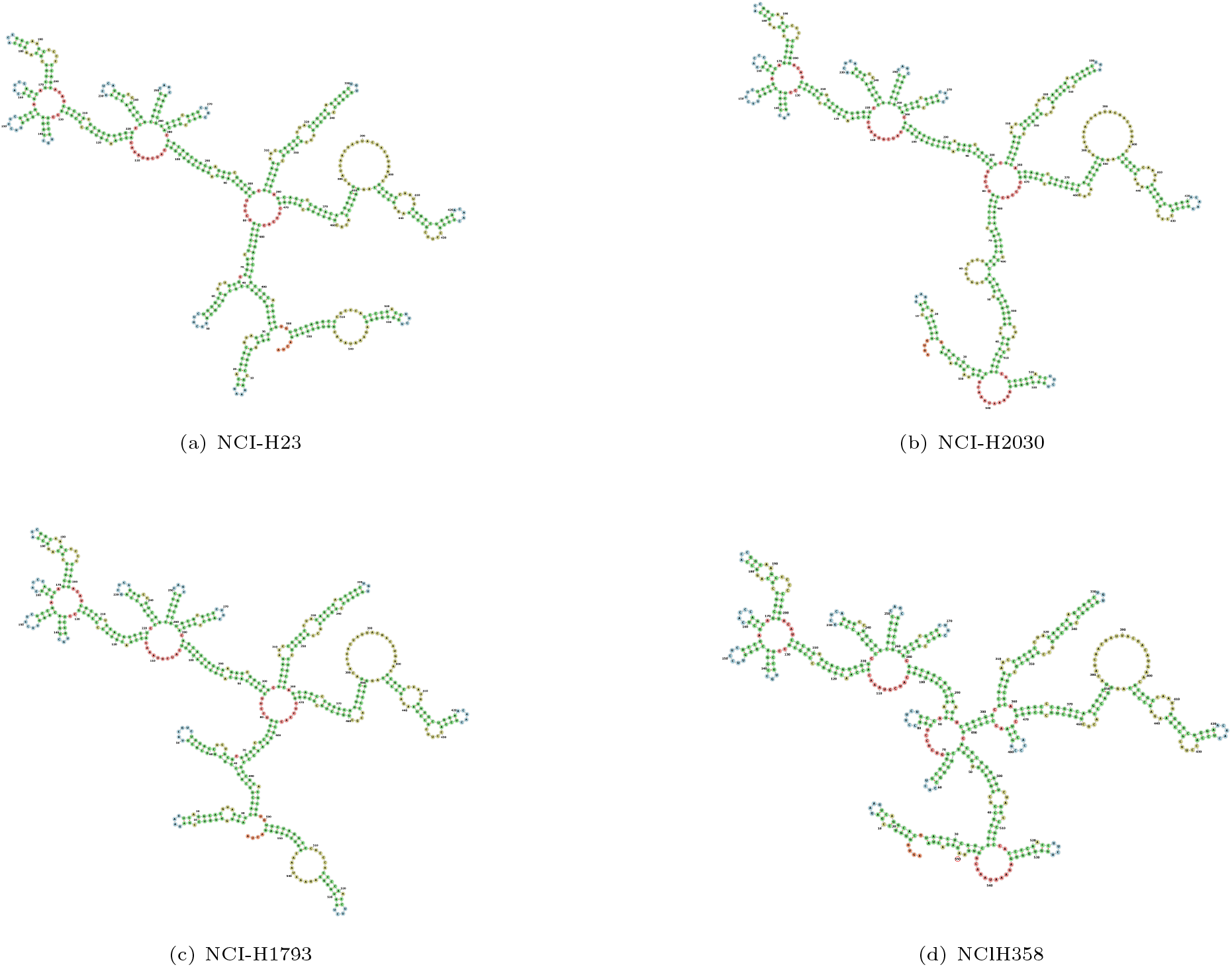
RNA secondary structures of the Macrophage Migration Inhibitory Factor (MIF) transcript across four lung cancer cell lines: (a)NCI-H23, (b)NCI-H2030, (c)NCI-H1793, and (d)NClH358.

**Fig. 5.**
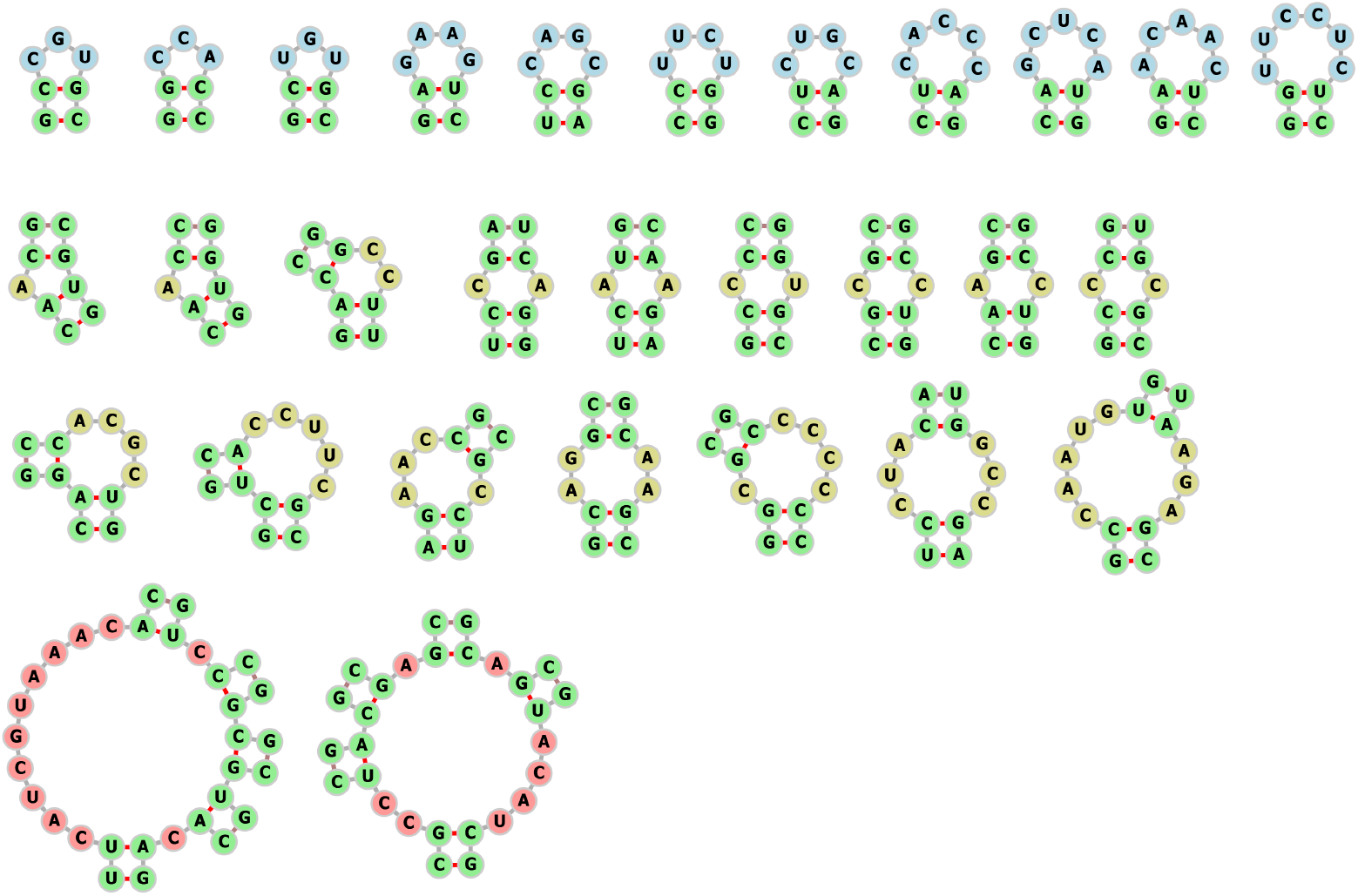
Conserved motifs identified across four lung cancer cell lines include hairpin loops, internal loops, and multibranch loops.

To further assess the druggability of these RNA motifs, we input them into the Inforna platform [13, 59], developed by the M. D. Disney research group. This platform integrates the world’s largest RNA-small molecule interaction database and employs machine learning algorithms to predict optimal binding ligands. Notably, the system successfully identifies multiple lead compounds with a perfect motif compatibility fitness score of 100; their specific structures and binding modes are shown in Fig. 3. Some of these molecules have already demonstrated promising RNA-binding activity in previous studies [60, 61]. These findings not only provide strong leads for the future development of MIF-targeting drugs but also open new avenues for expanding the application of RNA-targeted small molecules in cancer therapy.

## 3 Discussion

We propose a quantum algorithm for RNA motif identification that leverages advanced techniques in quantum computing. By encoding non-canonical base interactions into binary representations, we incorporate edge weights into the framework of the quantum subgraph isomorphism algorithm [62]. This approach enables the successful integration of RNA secondary structure graphs into a quantum circuit for efficient recognition and computation.

We conduct evaluations on two critical tasks using real datasets: identification of known motifs and discovery of unknown common motifs. Our algorithm is subsequently applied to systematically identify conserved motifs across four distinct lung cancer cell lines, following by the analysis of their potential as viable targets for small-molecule drugs. This advancement not only improves the efficiency of screening for RNA-targeting small molecules but also enhances their specificity, thereby offering a more precise and effective approach for the development of RNA-targeted therapeutics.

However, it should be emphasized that validating drug efficacy requires extensive wet-lab experimentation, potentially including the de novo design of novel molecular compounds. These essential but distinct research endeavors extend beyond the scope of the current investigation and have been intentionally reserved for subsequent dedicated studies.

Given the constraints of the quantum simulator, we restrict RNA secondary structure graphs to at most 16 nucleotides (with weight encoding). For longer RNA molecules, we partition them into multiple subgraphs and analyze independently. However, by employing the LH representation detailed in Section 4, a graph of *N* nucleotides is compactly encoded into a unitary matrix by using *q* = 2 log_2_ *N* + 3 qubits, which demonstrating the exponential quantum resource efficiency of our algorithm. Considering that the longest RNA chain is about 1 × 10^5^ nucleotides(less than 40 qubits in theory are needed), this approach holds significant promise for practical application under the qubit limitations of the NISQ era. Moreover, as the increasing of qubits number, multiple RNA secondary structure graphs can be encoded into the quantum circuit simultaneously. This capability facilitates efficient parallel computation for large-scale RNA analysis.

While the proposed quantum algorithm is successfully applied to RNA motif identification, some limitations remain, which we aim to address in future work. One key challenge is the issue of circuit depth. Encoding the adjacency matrix onto a quantum circuit entails a complexity of 𝒪(*N*^2^), which significantly constrains the algorithm’s scalability as the problem size increases. Another area of concern is the expressive capacity of permutation matrices. Further investigation is required to determine whether the permutation matrix ansatz sufficiently represent the permutation matrix *P* for large problem instances.

## 4 Methods

### Graph matching

Once we obtain secondary structure graphs of RNA and RNA motif annotated with base interactions, we transform the 3D structure identification task of RNA motifs into a 2D graph matching problem. Thus, identifying specific motifs within a single RNA structure and searching for the same motif across two RNA structures become instances of the subgraph isomorphism problem and the common subgraph problem, respectively. Both problems are known to be NP-hard.

### Graph isomorphism problem

For motif detection case, let us consider an RNA 2D structure represented by a graph *G* = {*V, E*}, in which a vertex *v* ∈ *V* represents a nucleotide and its base type, and an edge *e* ∈ *E* represents a base pairing or phosphodiester backbone interaction. Similarly, the 2D structure graph of an RNA or RNA motif can also be represented by a graph *G*_*m*_ = {*V*_*m*_, *E*_*m*_}, where the subscript ‘*m*’ denotes ‘a smaller RNA or a motif’. The problem is given a permutation relation (a permutation matrix *P*) between the graphs *G* and *G*_*m*_, identify their common substructures or determine whether the graph *G*_*m*_ is a subgraph of the graph *G*. Let the adjacency matrices of *G* and *G*_*m*_ be *A*_*R*_ and *A*_*m*_, respectively. Thus we have the loss function,

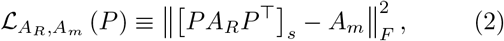

where [*A*]_*s*_ for an adjacency matrix *A* means a subblock of *A* with the same size as *A*_*m*_, and ∥·∥_*F*_ denotes the Frobenius norm. The decision variable in this optimization problem is the permutation matrix *P*, and the solution to the graph isomorphism problem is to find a *P* such that the loss function *L* is as small as possible.

### Quantum algorithms for graph isomorphism

Our work is inspired by Mariella and Simonetto of their quantum subgraph isomorphism algorithm [62] based on undirected unlabeled graphs. The principle of this algorithm and the extensions we have made are describe in following. For simplicity, hereinafter we do not consider the subgraph case, which can be generalized by performing a block selection operation on a bigger graph.

From above, we have the objective function and the decision variables for the graph isomorphism problem.

However, implementing Eq. (2) on a quantum circuit directly means that all *N*^2^ matrix elements of two *N* dimensional matrices need to be subtracted, which is obviously very inefficient. Fortunately, we have the following lemma:

#### Lemma 1.

For any *x, y* ∈ ℤ_2_, it holds that

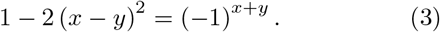

Moreover, we note the right-hand side of Eq. (3) can be further written as

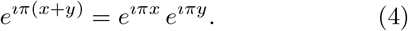

Thus, Lemma 1 together with Eq. (4) enable the conversion of the subtraction of each matrix element in Eq. (2) into a multiplication form. Specifically, for two *N* × *N* adjacency matrices *A* and *B*, with their matrix elements *A*_*i,j*_, *B*_*i,j*_ ∈ ℤ_2_, the difference between *A* and *B* can be measured by,

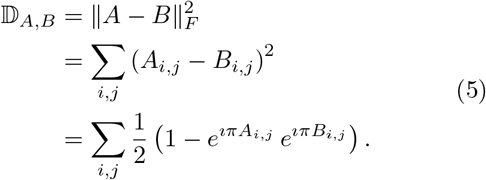

That is the reason that we binarized the adjacency matrix in the previous section.

Eq. (5) shows that 𝔻_*A,B*_ can be represented as a Hadamard product of two *N* × *N* matrices, which is still difficult to implement on quantum circuits. However, it can be re-represented as a trace of the product of two matrices if we put each element, for example 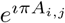, on the diagonal positions of an *N*^2^-dimensional zero matrix. Defining the unitary operator representation of the adjacency matrix,

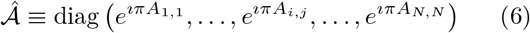

(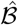 is similar for *B*, Eq. (5) can be re-represented as 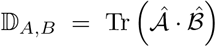. Here we have omitted irrelevant terms in Eq. (5) during RNA motif identification process. Since both 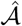 and 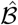 are diagonal matrices, their product is obviously diagonal. Thus, we can apply a state |*ψ*⟩ ∝ (1, 1, *· · ·*, 1)^*†*^ with all elements equal to 1 to achieve the trace, that is, 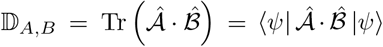. Noting that |+⟩^⊗*n*^ = *ψ*, given a quantum circuit with the initial state |0⟩^⊗*n*^, the state *ψ* can be prepared by applying Hadamard gates on each qubit. In this way, we obtain the **Log-Hadamard** representation of the adjacency matrice that can be encoded onto a quantum circuit to obtain the value of Eq. 5:

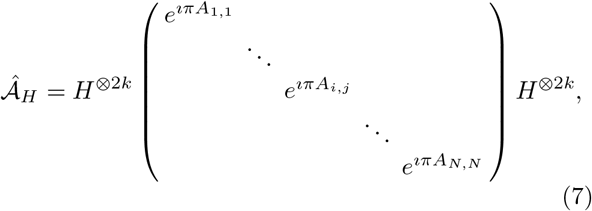

This can be implemented directly by the Diagonal_gate command in Qiskit.

Next, a variable permutation matrix *P* is employed to identify the isomorphism between the two graphs by minimizing 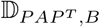. As the adjacency matrices are transfromed into diagonal from, the tensor product of permutation matrix *P* ⊗ *P* is then acts on 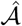 instead of *P*, which is easy to prove diag 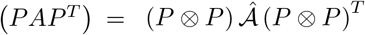. The permutation matrix *P* is represented by an approximate ansatz with parameters, based on the fact that (i) the product of permutation matrices is still a permutation matrix, and (ii) the permutation matrix is self-inverse. We parameterize the permutation matrix as

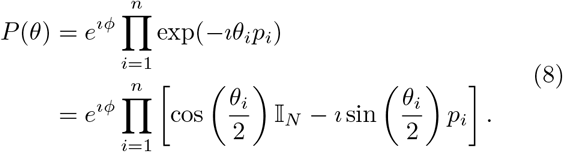

Note that |cos(*tπ/*2)| is equal to 1(0) when *t* is even (odd), and the case of |sin(*tπ/*2)| is exactly the opposite. Let each parameter in *θ* ≡{*θ*_1_, *θ*_2_, …, *θ*_*i*_} be an integer multiple of *π*. This allows the ansatz (8) to be expressed as a product of different combinations of *p*_*i*_. By constructing different forms of *p*_*i*_ such as *σ*_*x*_, 𝕀 ⊕ *σ*_*x*_, and etc., the approximate ansatz of the permutation matrix can be obtained.

Finally, we can use an ancillary qubit to construct Hadamard test to obtain the result. Note the expected value of the entire quantum circuit acting on the initial state |0⟩^⊗*n*^ is the negative of the loss function (2). The entire quantum circuit is presented in Fig. 1(b). For a graph with *N* vertices, 2 log_2_ *N* + 3 qubits are utilized, where *k* = log_2_ *N* and *k*^*′*^ = log_2_(*N*^*′*^). Here *N*^*′*^ represent the size of a smaller graph that used to determine isomorphism with the original graph. By configuring the Hadamard gate to operate exclusively on *k*^*′*^ + 1 qubits, the loss function can be adjusted to compute only the components relevant to the subgraph. In addition, an additional qubit is introduced in the permutation matrix to match the dimensions of the weighted adjacency matrix.

## Data Availability

The RNA motifs show in Fig. 1(c) and Fig. 2 are collected from RNA3DHud with version 3.87 (August.14, 2024) at https://www.bgsu.edu/research/rna/databases.html. And the 3D structure graphs of RNA chains show in Fig. 1(c) and (d) are downloaded from the PDB database at https://www.rcsb.org/.

## Code Availability

The code that supports the findings of this study is available from the authors upon reasonable request.

## Acknowledgments

We acknowledge support from State Key Laboratory of Genome and Multi-omics Technologies, China National GeneBank (CNGB) and the Guangdong Bigdata Engineering Technology Research Center for Life Sciences.

## Competing Interests

The authors declare no competing interests.

## Notes

### Competing Interest Statement

The authors have declared no competing interest.

